# Sex differences in gene expression in response to ischemia in the human myocardium

**DOI:** 10.1101/282350

**Authors:** Gregory Stone, Ashley Choi, Meritxell Oliva, Joshua Gorham, Mahyar Heydarpour, Christine E. Seidman, Jon G. Seidman, Sary F. Aranki, Simon C. Body, Vincent J. Carey, Benjamin A. Raby, Barbara E. Stranger, Jochen D. Muehlschlegel

## Abstract

**Background:** Sex differences exist in the prevalence, presentation, and outcomes of ischemic heart disease. Females have higher risk of heart failure post myocardial infarction relative to males and the female sex is an independent risk factor for hospital and operative mortality after cardiac surgery. However, the mechanisms underlying this sexual dimorphism remain unclear. We examined sex differences in human myocardial gene expression in response to ischemia.

**Methods:** Left ventricular biopsies from 68 male and 46 female patients undergoing aortic valve replacement surgery were obtained at baseline and after a median 74 minutes of cold cardioplegic arrest/ischemia and respective transcriptomes were quantified by RNA-Seq. Sex-specific responses to ischemia were quantified by differential gene expression, expression quantitative trait loci (eQTL) and pathway and functional analysis. Cell-type enrichment analysis. was used to obtain an estimate of the identity and relative proportions of different cell types present in each sample.

**Results:** A sex-specific response to ischemia was observed for 271 genes. Functional annotation analysis revealed sex-specific modulation of the oxytocin signaling pathway and common pathway of fibrin clot formation. The eQTL analysis identified variant-by-sex interaction eQTLs at baseline and post-ischemia, indicative of sex differences in the genotypic effects on gene expression, and cell-type enrichment analysis showed sex-bias in proportion of specific cell types.

**Conclusion:** In response to myocardial ischemia, the human left ventricle demonstrates changes in gene expression that differ between the sexes. These differences provide insight into the sexual dimorphism of ischemic heart disease and may aid in the development of sex-specific therapies that reduce myocardial injury.

## Introduction

There is a growing recognition of sex differences in ischemic heart disease outcomes. Women with non-obstructive coronary artery disease have a higher mortality and event rate than men,^1^ and are more likely to develop heart failure after myocardial infarction.^2^ The operative mortality of females undergoing coronary artery bypass grafting (CABG) is also higher than for men.^3^ Despite the systematic differences between men and women in the presentation and outcomes of ischemic heart disease, women remain grossly underrepresented in research studies, and sex-specific outcomes are spuriously reported.^4^ We have previously investigated changes in gene expression due to ischemia in the human left ventricle (LV) using cardioplegia and cardiopulmonary bypass (CPB) as a reproducible human ischemic model of myocardial infarction.^5^ While there has been some investigation into sex-specific transcription in the healthy myocardium,^6^ a dearth of knowledge remains regarding sex differences in the transcriptional response to myocardial ischemia. To date, most sex-specific investigations have focused on sex hormones and genes on the sex chromosomes.^7^ However, evidence exists to suggest that genes on autosomal chromosomes may show sex differences in the response to ischemia, and that this variable response has implications for sex-specific outcomes.^8^

We employed whole-genome RNA sequencing to characterize sex differences in the human transcriptional response to myocardial ischemia and to identify sex-biased genomic loci that contribute to variation in expression levels of RNA after ischemia, namely expression quantitative trait loci (eQTLs). To our knowledge, this is the first study investigating sex differences in ischemic heart disease at the transcriptional level.

## Methods

### Patients and Tissue Samples

118 patients undergoing elective aortic valve replacement surgery were prospectively enrolled after obtaining written informed consent approved by the Partners Healthcare Institutional Review Board (Boston, MA) (Table 1). Two punch biopsies from the anterolateral apical LV wall, one pre-(baseline) and one post-ischemia, were obtained. Baseline samples were collected immediately after commencement of CPB, and post-ischemia samples were obtained after a median of 74 min of ischemic time (range: 41-195 min) that included intermittent cold blood cardioplegia for myocardial protection.

**Table 1.**
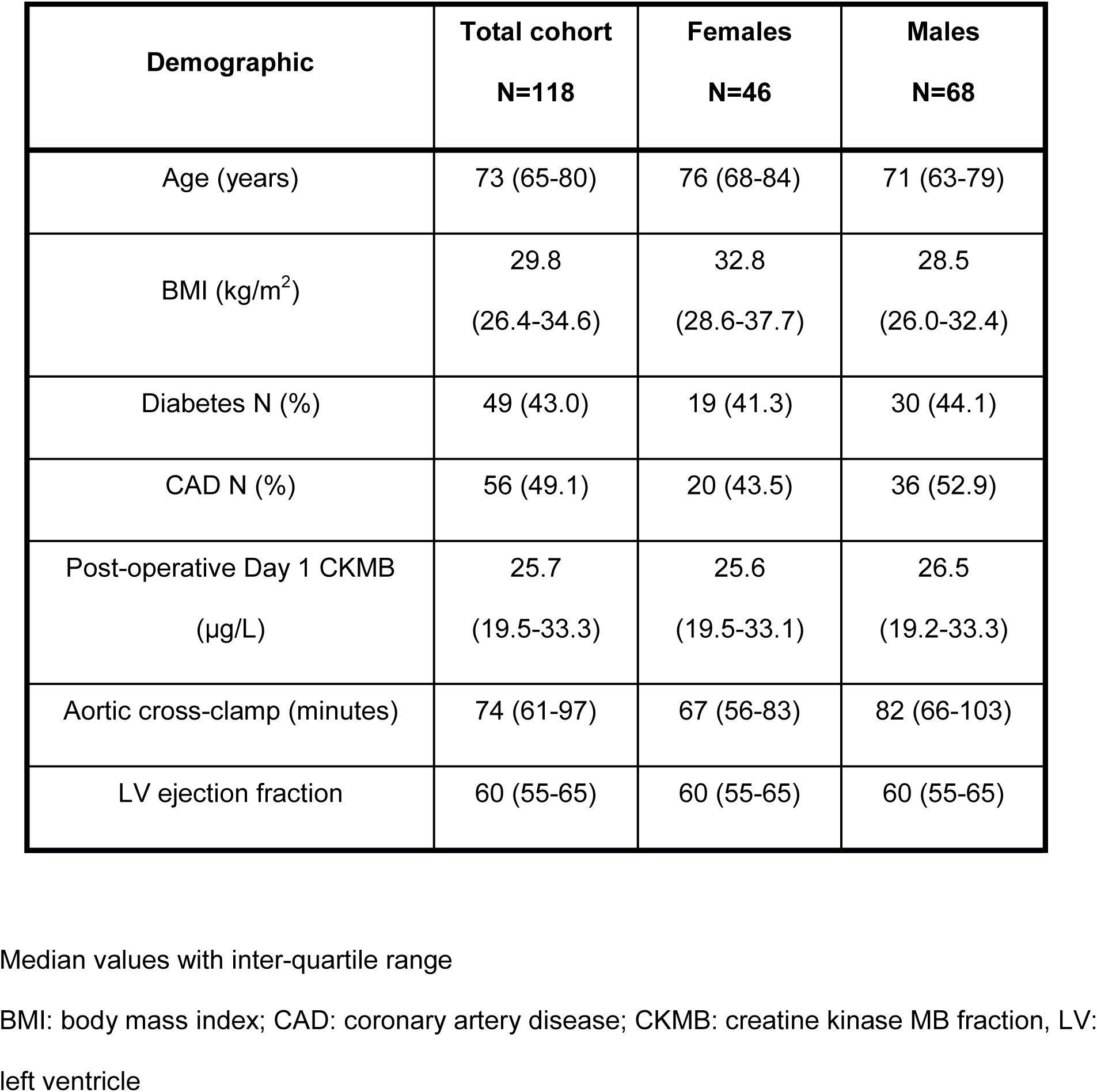
Patient demographics and clinical characteristics

| **Demographic** | **Total cohort**<br>**N=118** | **Females**<br>**N=46** | **Males**<br>**N=68** |
| --- | --- | --- | --- |
| Age (years) | 73 (65-80) | 76 (68-84) | 71 (63-79) |
| BMI (kg/m<sup>2</sup>) | 29.8<br>(26.4-34.6) | 32.8<br>(28.6-37.7) | 28.5<br>(26.0-32.4) |
| Diabetes N (%) | 49 (43.0) | 19 (41.3) | 30 (44.1) |
| CAD N (%) | 56 (49.1) | 20 (43.5) | 36 (52.9) |
| Post-operative Day 1 CKMB<br>(µg/L) | 25.7<br>(19.5-33.3) | 25.6<br>(19.5-33.1) | 26.5<br>(19.2-33.3) |
| Aortic cross-clamp (minutes) | 74 (61-97) | 67 (56-83) | 82 (66-103) |
| LV ejection fraction | 60 (55-65) | 60 (55-65) | 60 (55-65) |

Median values with inter-quartile range
BMI: body mass index; CAD: coronary artery disease; CKMB: creatine kinase MB fraction, LV: left ventricle

### RNA sequencing, alignment, and annotation

RNA sequencing and preservation techniques have been detailed elsewhere.^55^ Briefly, tissue samples were preserved in RNAlater® (Ambion, Life Technologies, USA) immediately after collection and sequenced in batches at a later time-point on the Illumina HiSeq 2000 (Illumina, San Diego, CA). Paired-end reads 50, 90, or 100 base pairs long were generated, producing ~134 million reads per sample. Adapter sequences were removed using Skewer,^9^ and reads were trimmed using Sickle^10^ at a quality threshold of 5. Remaining reads were aligned to the UCSC hg19 reference human genome using STAR (v2.5.2b).^11^ Aligned reads were subsequently counted using featureCounts (v1.5.1).^12^ Gene counts were pre-processed by applying a variance-stabilizing transformation and count normalization^13^. Biological sex was verified by measuring the baseline expression levels of *XIST*, an X-linked gene expressed significantly higher in females than in males at baseline.^6^ Four samples were removed due to inconsistencies in *XIST* expression.

### Differential Expression Analysis

We consider two forms of differential expression: 1) difference in expression level in response to ischemia in each sex (e.g. in the post-ischemia sample, gene A is expressed higher in females than in males), and 2) differential response to ischemia by sex (e.g. gene A increases it’s expression from baseline to post-ischemia in females, but not in males).

All statistical analyses were performed in R (v3.3.1). Read counts of genes were normalized while incorporating sample-level weights using voom,^14^ and analyzed for differential expression with limma (v3.30.11).^15^ Genes were filtered for inclusion (e.g. low abundance reads, low count across all samples) in accordance with practices established by the authors of edgeR.^16^ A linear model was constructed to calculate the mean expression of a gene, denoted y, in each sex at baseline and at post-ischemia while adjusting for age, diabetes, and ischemic time:

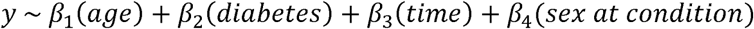

where denotes regression coefficients, and the factor “sex at condition” has 4 levels: female expression at baseline, female expression at post-ischemia, male expression at baseline, and male expression at post-ischemia. Subject is treated as a random effect with the limma *duplicateCorrelation* function.

1. To identify genes with significant **difference in expression level in response to ischemia in each sex**, the significance of the following model-fitted contrasts were assessed:

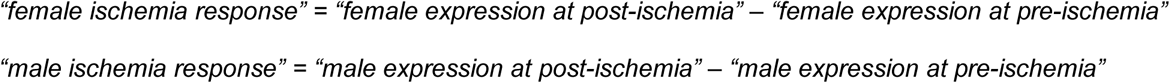
2. To determine if the **change in the expression of a gene in response to ischemia differed between the sexes**, the model was fit to the following contrast, which produced an estimated log_2_ fold change that measured the difference between the change in gene expression within females and that within males in response to ischemia:

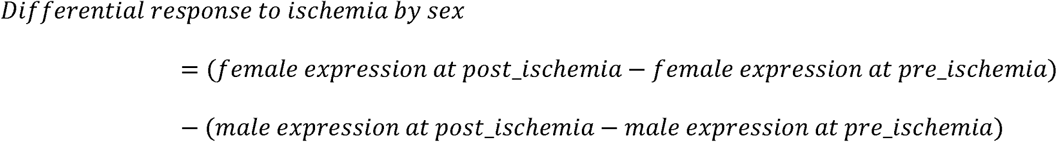

The significance (p_adj_ < 0.05) of each contrast was determined by empirical Bayes moderated t-statistics and adjustment for multiple testing was performed using Benjamini-Hochberg false discovery rate.^17^ A **positive**/**negative** log_2_ fold change is interpreted as an **increase**/**decrease** in gene expression due to ischemia in females relative to that in males. A significant “*Differential response to ischemia by sex”* contrast is compatible with multiple scenarios (see Figure 1 for illustrations):

**Figure 1.**
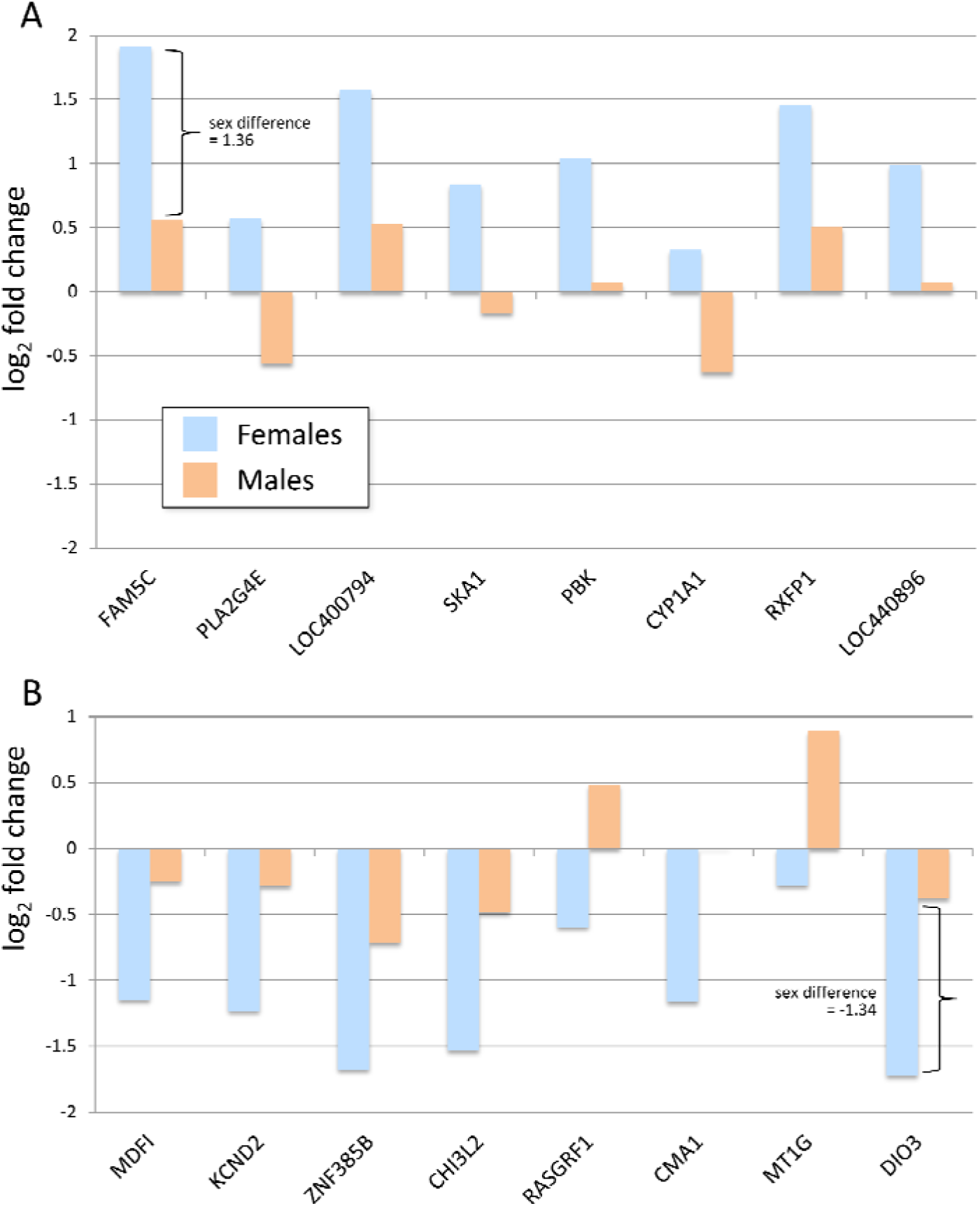
Title: Multidimensional scaling plot showing separation of samples by sex. The distance between any pair of samples is the log_2_ fold change between the root-mean-square deviation of counts of the top 500 genes. A shorter distance indicates a greater similarity between a pair of samples. X and Y-axis show distances in the 3^rd^ and 4^th^ dimension, respectively. Male samples are coral and female samples are turquoise.

1. Gene expression significantly increases/decreases, in a direction **opposite** of the other sex. 2) Gene expression significantly increases/decreases in both sexes, but more significantly in one sex. 3) Gene expression significantly increases/decreases only in one sex.

### Functional Analysis of Differentially Expressed Genes

Functional analysis of the genes with sex-specific differential expression was performed using g: Profiler^18^, which searches functional terms from GO, KEGG, REACTOME, CORUM, miRBase, and Human Phenotype Ontology databases. Significant functional terms were identified at Benjamini-Hochberg false discovery rate (FDR) =<10%.^17^ GO and KEGG associations elucidated genes involved in biological pathways. REACTOME enrichments identified genes involved specific biological processes and reactions, and CORUM associations revealed genes implicated in protein complexes. miRBase terms denoted miRNAs thought to target associated genes, and TRANSFAC terms indicated transcription factor binding sites possessed by listed genes. Human Phenotype Ontology (HP) enrichments revealed genes associated with a specific human phenotype.

### Expression quantitative trait loci (eQTL)

Using DNA isolated from whole blood, SNP genotyping was performed using the Illumina Omni2.5 with exome content genotyping array (Illumina, San Diego, CA). SNPs were excluded if the minor allele frequency (MAF) was less than 5%, the genotype missing rate was greater than 5%, or were not in Hardy-Weinberg equilibrium (P<10^−6^). These filters were applied to the non-imputed genotypes of males and females separately, with only the SNPs surviving in both groups retained for testing. The associations between SNP genotypes and gene expression were tested using Matrix eQTL, separately for baseline and post-ischemia profiles. We inferred hidden factors affecting gene expression for both sexes combined and both sexes independently using Probabilistic Estimation of Expression Residuals (PEER).^19^ A cis-eQTL analyses (within 100,000 base pairs from the transcription start site) was performed to identify and compare cis-eQTLs present at baseline and post-ischemia. To identify eQTL with different effects sizes in males and females at post-ischemia, an interaction cis-eQTL analysis was performed using matrix eQTL.^13^ A linear model was constructed to test the interaction of sex and SNP genotype on PEER-corrected residuals while adjusting for age, ischemic time, diabetes, sequence center, number of reads, and genetic principal components (PC1, PC2 and PC3).^13^ A negative interaction term indicates a larger eQTL effect size in females, whereas a positive value indicates a larger eQTL effect size in males. We performed this analysis separately for samples collected at baseline and at post-ischemia. We retained those genes with a significant SNP-by-sex interaction eQTL (Benjamini-Hochberg FDR=5%) only in the post-ischemia analysis for downstream analysis. Multiple-testing adjustment was done for all cis-eQTL analyses: EigenMT^20^ to correct for the number of SNPs within genes, and Benjamini-Hochberg to correct for the number of genes.

To test the shared effects between differentially expressed genes due to ischemia and the differential response to ischemia by sex, we quantified the proportion of true positives estimated from the enrichment of significant *P*-values (pi1).^21^

### Cell-type deconvolution

The tissue biopsy and resulting transcriptome profile are not restricted to a single cell-type. We attempted single-cell RNA sequencing using two different techniques (Fluidigm C1 and 10x Genomics Chromium) but were not successful given the size of human myocytes. Therefore, we performed a cell-type enrichment analysis to obtain an estimate of the identity and relative proportions of different cell types present in each sample, and to compare them to other tissues including heart left ventricle characterized by the Genotype-Tissue Expression (GTEx) Project. The data used for the analyses described in this manuscript were obtained from the GTEx Portal on 12/1/2017.^22^

We performed cell-type enrichment analysis using xCell^23^ on the transcriptomes of all11,688 GTEx v7 samples and the pre- and post-ischemia left ventricle biopsies. Sample enrichment scores were obtained for 64 immune and non-immune cell types. For all cell types, we tested for sex differences in a) cell abundances in pre-ischemic samples and b) the ratio of cell abundances in post-versus pre-ischemia.

We applied Principal Component Analysis (PCA) to the matrix of cell types and sample enrichment scores to ensure that the observed differences in expression between females and males are not a result of variable expression across heterogeneous cell types. Furthermore, we explored the contribution of each cell type to differential gene expression between sexes in response to ischemia.

## Results

We define up-regulated and down-regulated genes as those genes whose expression differs in the pre- and post-ischemic states (irrespectively of sex), where up-regulation refers to an increase in gene expression from the pre- to post-ischemia state and down-regulation to a decrease. We define sex-biased response genes as those that demonstrate significant sex differences in either the magnitude or the direction of the post-ischemia response. Notice that the sex-bias definition is exclusively restricted to the response to ischemia rather than to transcriptomic sex differences at baseline.

### Patient Demographics

Of the 114 patients in this study, 68 (60%) are male and 46 (40%) are female. The median age is 73 years, with all women being postmenopausal.

### Gene expression

66 million paired transcripts per sample were available for alignment after adapter and quality trimming, and 84.5% of reads were uniquely aligned to the human genome (UCSC hg19) with STAR.

The filter threshold for genes to be included in differential expression analysis was set to 3 counts per million (gene counts x 1,000,000 ÷ library size) in at least 2 samples (Supplemental Figure 1 shows the effect of filtering on stabilizing mean-variance trend). 15,265 genes were analyzed for differential expression in response to ischemia: 8,417 were differentially expressed in both sexes, 1,263 genes were differentially expressed in females only, and 2,266 genes were differentially expressed in males only (Supplemental Table 1). For 271 genes, we observed a *differential response to ischemia by sex* (as determined by formal interaction testing, p_adj_ < 0.05) (Supplemental Table 2) – 70 genes changed in only males, 96 changed in only females, and 105 changed in both sexes but to different degrees. Several of these genes, such as Estrogen Receptor 1 (ESR1), centromere protein L (CENPL), secreted phosphoprotein 1 (SPP1) have also been shown to express differently been males and females in previous studies,^24, 25^ validating our results.

The effect of sex on gene expression was visualized with a multidimensional scaling plot, which revealed stratification by sex in the 3^rd^ dimension, showing a clear demarcation between males and females (Figure 2). Furthermore, in the cluster plot in Figure 3, we show the separation of genes with significant change that present in only females, in only males, and in females and males combined.

**Figure 2.**
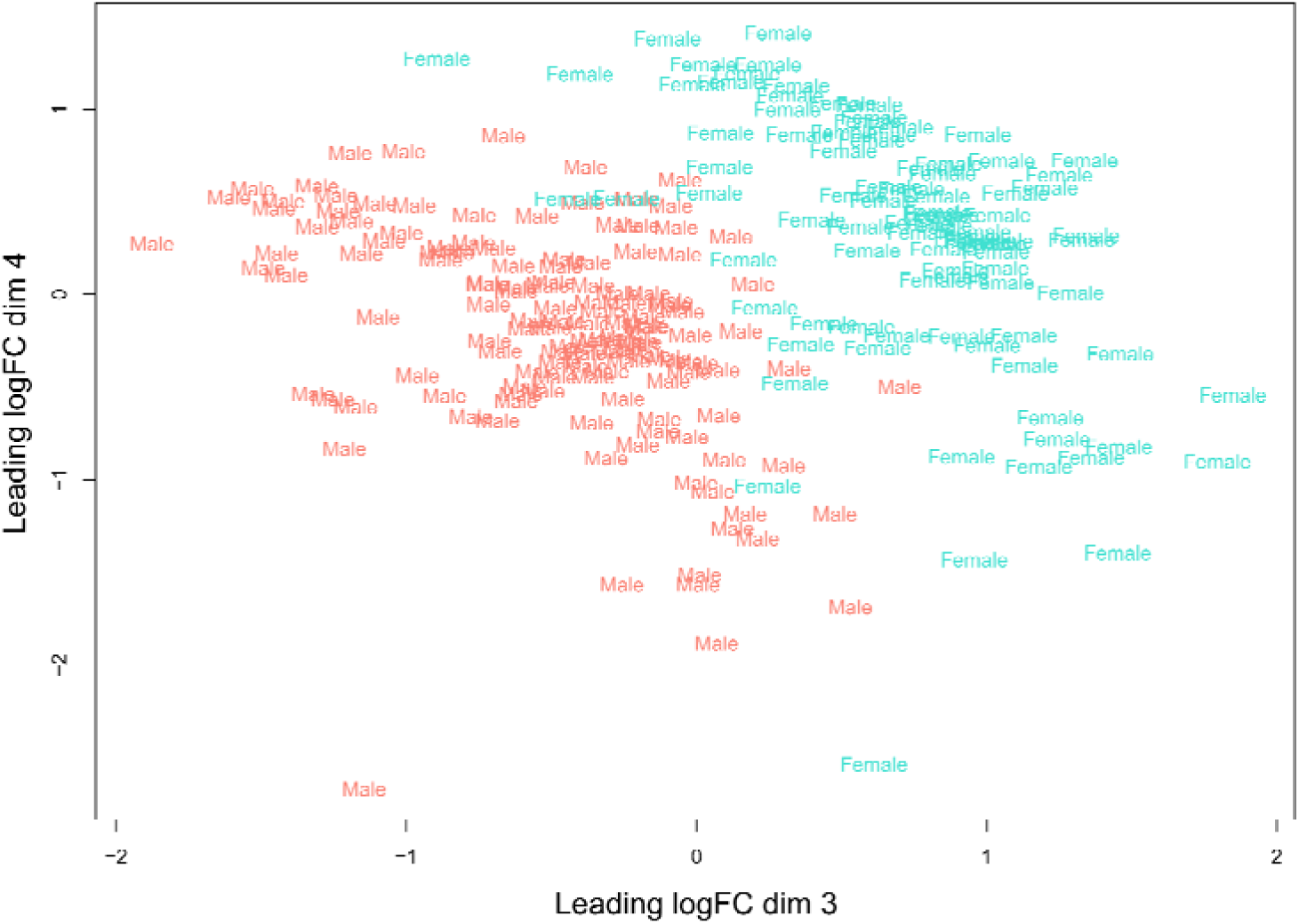
Title: Relative change in gene expression due to ischemia. Orange and blue bars represent the mean log_2_ fold change in gene expression in female and male samples, respectively, due to ischemia. The black brackets denote examples of the changes in gene expression in females relative to males due to ischemia.

**Figure 3:**
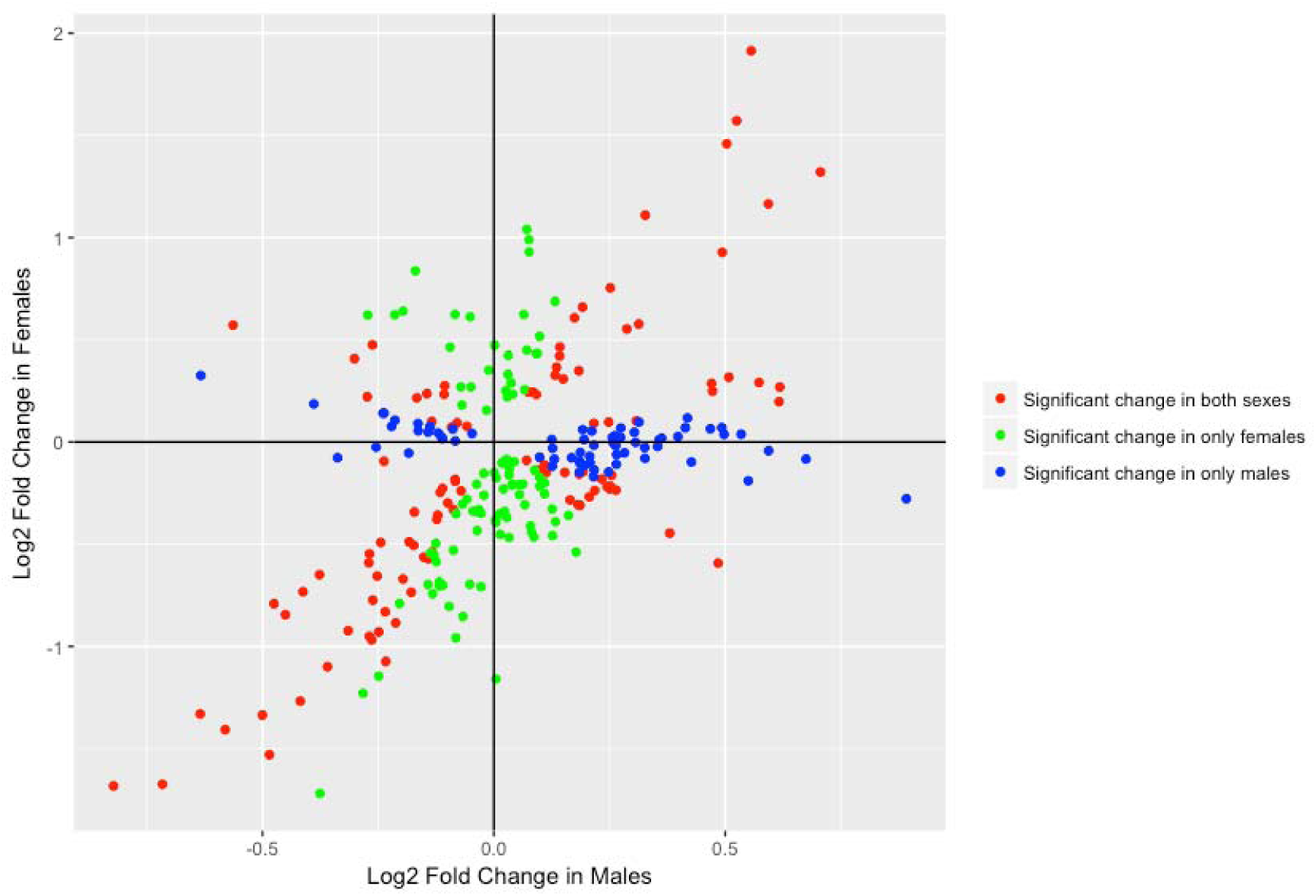
Title: DNA sequence variant rs11985898 has a sex-specific effect on the expression of *LETM2* at only Baseline. The boxplots in pink and blue show female and male *LETM2* expression, respectively. The Y-axis denotes expression levels, whereas the X-axis shows, from left to right, homozygous for the reference allele, heterozygous, and homozygous for the alternate allele genotypes with the number of corresponding samples.

### Relative up-regulation in females

The gene with the largest difference between females and males in transcriptional response to ischemia was *FAM5C*, which was up-regulated in response to ischemia in both males and females, but to a significantly greater degree in females (Table 2). We also observed a sex difference in expression response to ischemia for *PLA2G4E*, which encodes the protein phospholipase A_2_ Group IVE (cPLA_2_E). In response to ischemia, PLA2G4E was up-regulated in females, but down-regulated in males. Another gene with a differential response to ischemia by sex was cytochrome P450-family 1-subfamily A-polypeptide 1 (*CYP1A1)*, which was down-regulated in males and up-regulated in females in response to ischemia.

**Table 2.**
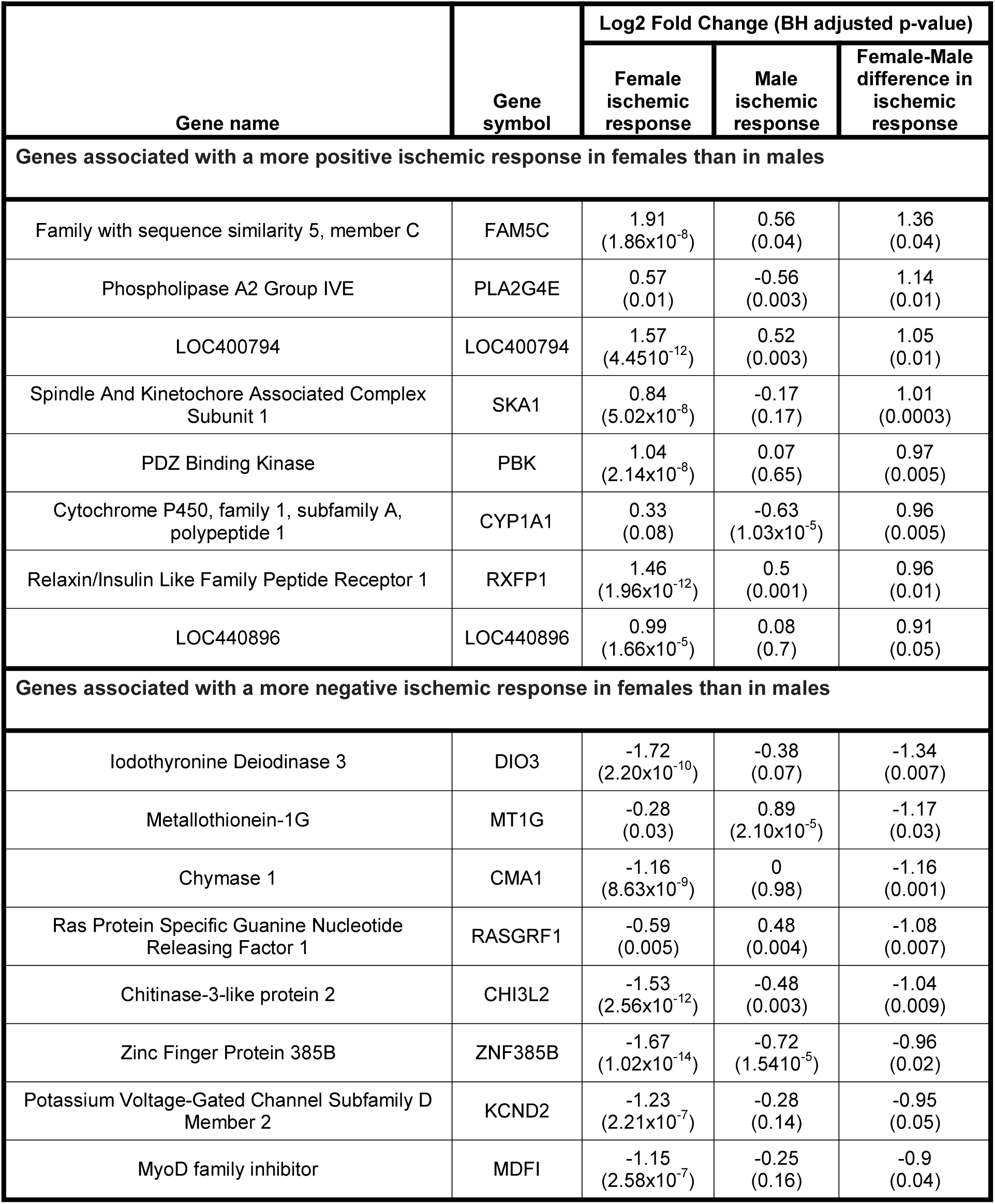
Top 8 up- and down-regulated genes in females that are significantly differently regulated relative to males.

| Gene name | Gene symbol | Log2 Fold Change (BH adjusted p-value) |  |  |
| --- | --- | --- | --- | --- |
|  |  | Female ischemic response | Male ischemic response | Female-Male difference in ischemic response |
| Genes associated with a more positive ischemic response in females than in males |  |  |  |  |
| Family with sequence similarity 5, member C | FAM5C | 1.91<br>(1.86x10<sup>-8</sup>) | 0.56<br>(0.04) | 1.36<br>(0.04) |
| Phospholipase A2 Group IVE | PLA2G4E | 0.57<br>(0.01) | -0.56<br>(0.003) | 1.14<br>(0.01) |
| LOC400794 | LOC400794 | 1.57<br>(4.4510<sup>-12</sup>) | 0.52<br>(0.003) | 1.05<br>(0.01) |
| Spindle And Kinetochore Associated Complex Subunit 1 | SKA1 | 0.84<br>(5.02x10<sup>-8</sup>) | -0.17<br>(0.17) | 1.01<br>(0.0003) |
| PDZ Binding Kinase | PBK | 1.04<br>(2.14x10<sup>-8</sup>) | 0.07<br>(0.65) | 0.97<br>(0.005) |
| Cytochrome P450, family 1, subfamily A, polypeptide 1 | CYP1A1 | 0.33<br>(0.08) | -0.63<br>(1.03x10<sup>-5</sup>) | 0.96<br>(0.005) |
| Relaxin/Insulin Like Family Peptide Receptor 1 | RXFP1 | 1.46<br>(1.96x10<sup>-12</sup>) | 0.5<br>(0.001) | 0.96<br>(0.01) |
| LOC440896 | LOC440896 | 0.99<br>(1.66x10<sup>-5</sup>) | 0.08<br>(0.7) | 0.91<br>(0.05) |
| Genes associated with a more negative ischemic response in females than in males |  |  |  |  |
| Iodothyronine Deiodinase 3 | DIO3 | -1.72<br>(2.20x10<sup>-10</sup>) | -0.38<br>(0.07) | -1.34<br>(0.007) |
| Metallothionein-1G | MT1G | -0.28<br>(0.03) | 0.89<br>(2.10x10<sup>-5</sup>) | -1.17<br>(0.03) |
| Chymase 1 | CMA1 | -1.16<br>(8.63x10<sup>-9</sup>) | 0<br>(0.98) | -1.16<br>(0.001) |
| Ras Protein Specific Guanine Nucleotide Releasing Factor 1 | RASGRF1 | -0.59<br>(0.005) | 0.48<br>(0.004) | -1.08<br>(0.007) |
| Chitinase-3-like protein 2 | CHI3L2 | -1.53<br>(2.56x10<sup>-12</sup>) | -0.48<br>(0.003) | -1.04<br>(0.009) |
| Zinc Finger Protein 385B | ZNF385B | -1.67<br>(1.02x10<sup>-14</sup>) | -0.72<br>(1.5410<sup>-5</sup>) | -0.96<br>(0.02) |
| Potassium Voltage-Gated Channel Subfamily D Member 2 | KCND2 | -1.23<br>(2.21x10<sup>-7</sup>) | -0.28<br>(0.14) | -0.95<br>(0.05) |
| MyoD family inhibitor | MDFI | -1.15<br>(2.58x10<sup>-7</sup>) | -0.25<br>(0.16) | -0.9<br>(0.04) |

### Relative down-regulation in females

The gene with the largest sex difference in response to ischemia is *DIO3*, which encodes the protein iodothyronine deiodinase 3 (D3) (Table 2). *DIO3* expression was down-regulated in both sexes in response to ischemia, but to a significantly greater degree in females. Additionally, *MT1G*, which encodes the protein Metallothionein-1G, was significantly down-regulated in females in response to ischemia and did not significantly change in males. In mice, metallothionein protects against ischemic injury.^53^ Also, *CMA1*, which encodes the protein chymase, was significantly down-regulated in females and showed no change in males in response to ischemia.

### Functional Annotation

Functional annotation analysis of the 271 genes that showed a sex-bias in response to ischemia revealed significant enrichment of several GO, KEGG, REACTOME, HP, CORUM, miRBase, and TRANSFAC terms (Table 3, Supplemental Table 3). Genes in the oxytocin signaling pathway were enriched, which supports previous studies showing that oxytocin has been shown to be cardioprotective post infarction^26^ and after ischemia/reperfusion injury.^27^ Related genes were either down-regulated in response to ischemia in females and up-regulated in males, or down-regulated in both sexes, but to a greater degree in females. *PLCB3* was the only gene to be up-regulated in both sexes, but to a lesser extent in females. Additionally, though transcription of oxytocin receptor (*OXTR*) is down-regulated in the rat infarcted LV,^26^ *OXTR* was down-regulated in females while up-regulated in males. The female ischemic myocardium may therefore suppress the oxytocin signaling pathway more so than in males, thereby negating its protective effects and possibly inducing a vulnerability.

**Table 3.**
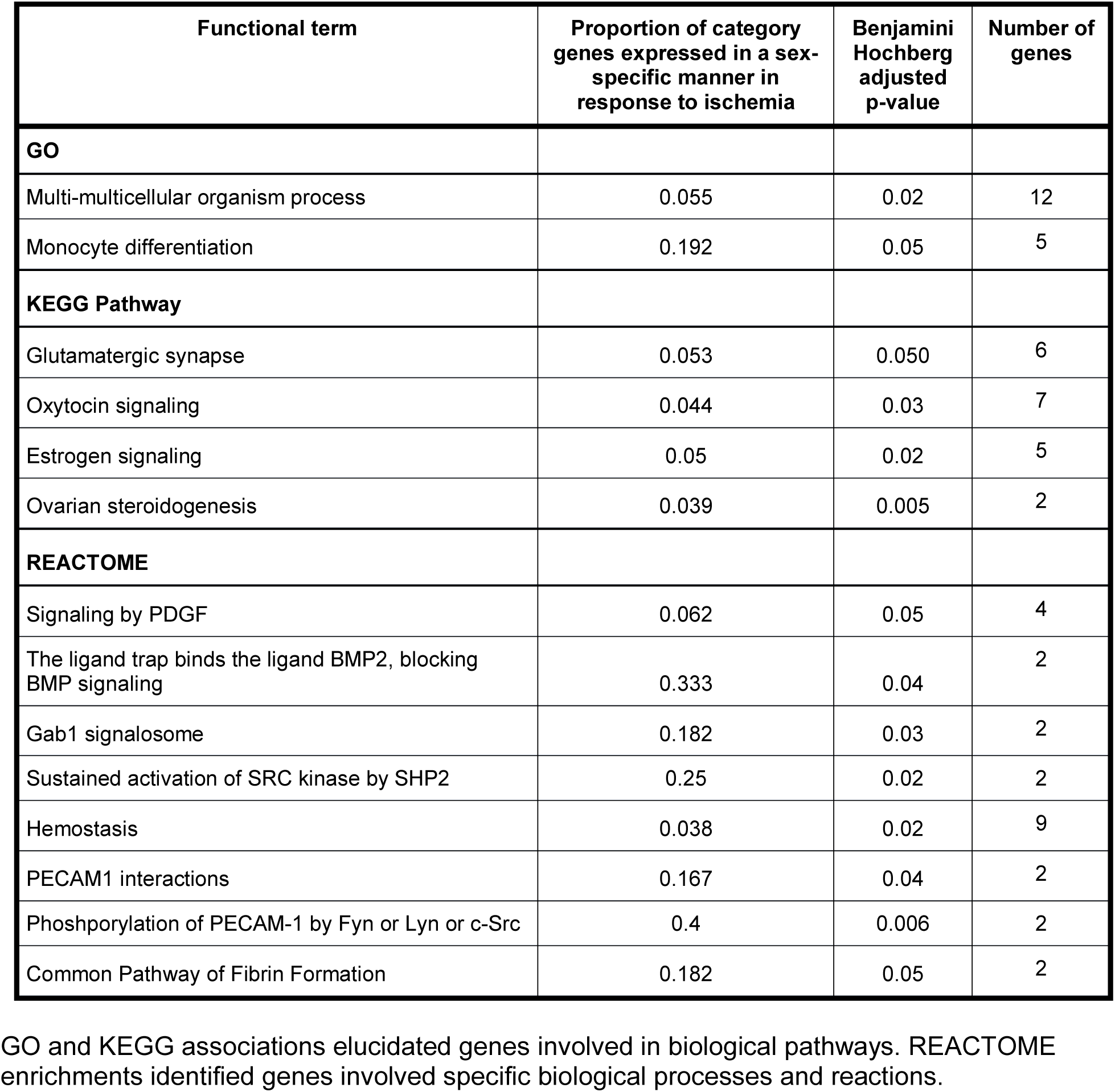
Functional analysis

| Functional term | Proportion of category genes expressed in a sex-specific manner in response to ischemia | Benjamini Hochberg adjusted p-value | Number of genes |
| --- | --- | --- | --- |
| **GO** |  |  |  |
| Multi-multicellular organism process | 0.055 | 0.02 | 12 |
| Monocyte differentiation | 0.192 | 0.05 | 5 |
| **KEGG Pathway** |  |  |  |
| Glutamatergic synapse | 0.053 | 0.050 | 6 |
| Oxytocin signaling | 0.044 | 0.03 | 7 |
| Estrogen signaling | 0.05 | 0.02 | 5 |
| Ovarian steroidogenesis | 0.039 | 0.005 | 2 |
| **REACTOME** |  |  |  |
| Signaling by PDGF | 0.062 | 0.05 | 4 |
| The ligand trap binds the ligand BMP2, blocking BMP signaling | 0.333 | 0.04 | 2 |
| Gab1 signalosome | 0.182 | 0.03 | 2 |
| Sustained activation of SRC kinase by SHP2 | 0.25 | 0.02 | 2 |
| Hemostasis | 0.038 | 0.02 | 9 |
| PECAM1 interactions | 0.167 | 0.04 | 2 |
| Phosphorylation of PECAM-1 by Fyn or Lyn or c-Src | 0.4 | 0.006 | 2 |
| Common Pathway of Fibrin Formation | 0.182 | 0.05 | 2 |

GO and KEGG associations elucidated genes involved in biological pathways. REACTOME enrichments identified genes involved specific biological processes and reactions.

Sex-biased genes were also enriched in the common pathway in fibrin formation and hemostasis. The coagulation factor X (*F10*) was up-regulated to a greater degree in males than females while thrombomodulin (*THBD*) was down-regulated to a greater degree in females than males in response to ischemia. *F10* and *THBD* expression have both been shown to be cardioprotective in animal models.^28, 29^ Additionally, *SERPINA5*, which encodes Protein C inhibitor that inhibits thrombin-thrombomodulin complex formation,^30^ was down-regulated in both sexes in response to ischemia, but to a greater magnitude in females. Sex-specific changes in hemostasis due to myocardial ischemia, and the mediation of hemostasis by the LV, require further investigation.

### eQTL identification

We performed eQTL analysis to assess whether genetic variation influences sex differences in ischemic response. Of the 114 patients in this study, complete genotype information was obtained for 110 (66 males, 44 females). After applying filtering criteria, 663,575 SNPs were tested for eQTLs, with 14,967 genes at baseline and 15,251 genes at post-ischemia. Cis-eQTL analyses at baseline and post-ischemia were performed separately: 1) females only, 2) males only and 3) females and males combined. cis-eQTLs at baseline for 95 genes were identified exclusively in females, 359 genes exclusively in males, and 932 genes in females and males combined (p_adj_ <= 0.05). Fewer post-ischemia cis-eQTLs were identified: 46 genes in females only, 317 genes in males only, and 804 genes in females and males combined (p_adj_ < 0.05). We identified SNP-by-sex interaction cis-eQTLs at baseline (*LETM2*, rs11985898, p_adj_ = 0.02), and at post-ischemia (*SLC30A7*, kgp2665681, p_adj_ = 0.04 and *VGLL4*, rs11707097, p_adj_ = 0.05), indicative of sex differences in the genotypic effects on gene expression (Figure 4, Supplemental Table 4) Out of the 271 genes that had gender-specific ischemic response, we have identified 7 genes that are significant eQTL at post-ischemia for females and males combined: ATP6AP1L, C1QTNF9, GSTM5, SPATA7, THEM178, WBSCR27 and ZNF385B.

**Figure 4:**
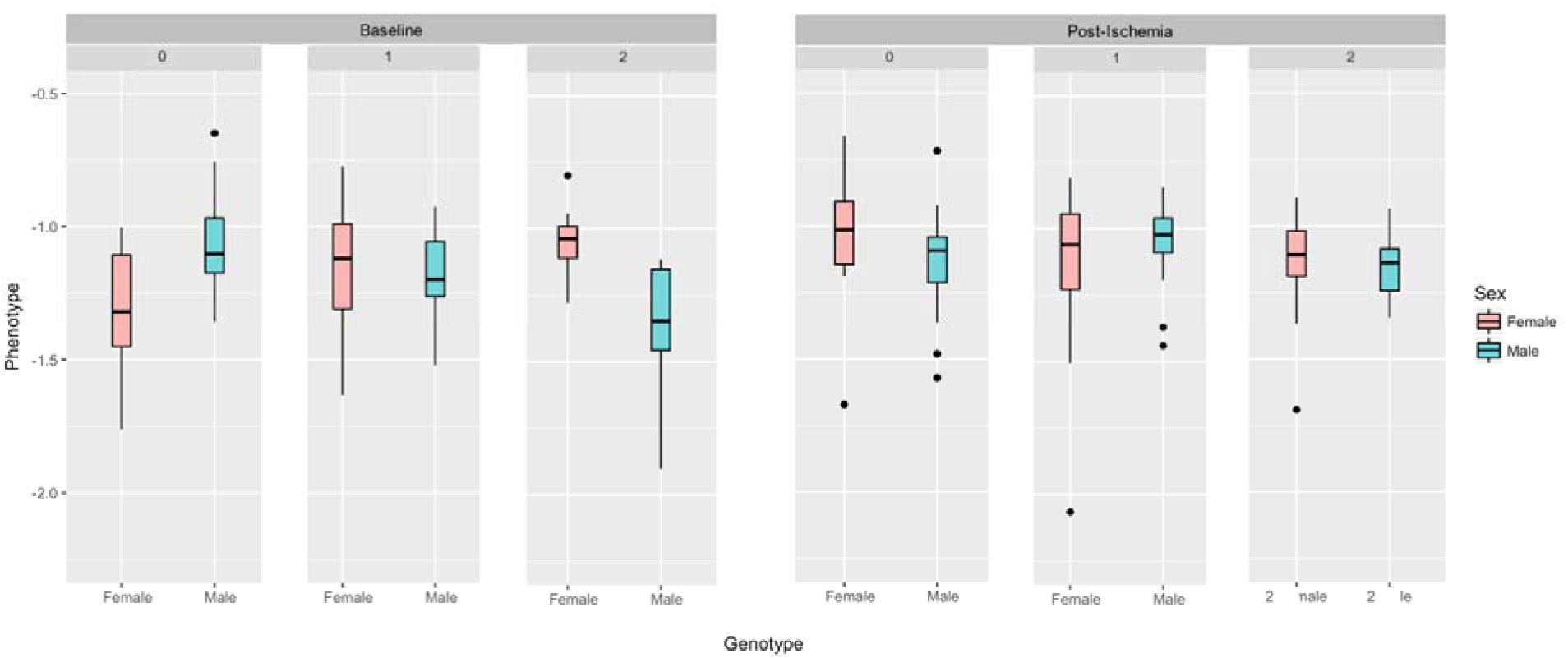
Example of an eQTL for gene *LETM2* DNA sequence variant rs11985898 has a sex-specific effect on the expression of *LETM2* at only Baseline. The boxplots in pink and blue show female and male *LETM2* expression, respectively. The Y-axis denotes expression levels, whereas the X-axis shows, from left to right, homozygous for the reference allele (0), heterozygous (1), and homozygous for the alternate allele (2) genotypes.

### Cell-type deconvolution

Clustering of the cell-type enrichment profiles of 220 samples revealed clustering by individual, and not by ischemia (Supplemental Figure 2a and b), demonstrating that intra-individual samples exhibit similar cell-type composition profiles that are not altered by ischemia.

Furthermore, when clustered with the multi-tissue GTEx cell-type enrichment profiles, the biopsy samples cluster with the GTEx left ventricular tissues, showing that the tissue collected is comparable between studies (Supplemental Figure 3).

At both baseline and ischemia conditions, we observe sex differences in cell-type enrichment scores (Supplemental Figure 4). Baseline samples exhibited significant sex differences in enrichment of several cell-types: hematopoietic stem cells, chondrocytes, macrophages M1 and immature dendritic cells are significantly more abundant in females than males (1.1, 1.22, 1.39, 1.47 female/male ratio, respectively; t-test P< 0.01). In female samples, we observed an increase in the abundance of common lymphoid progenitor cells post-ischemia relative to baseline (fold change 1.13), and a decrease in macrophages M2 abundance post-ischemia (fold change 0.89). These effects are not observed in males.

## Discussion

The female sex is an independent risk factor for hospital and operative mortality after cardiac surgery.^3^ Furthermore, females have higher risk of heart failure post-MI relative to males.^2^ While the transcriptional response to ischemia in the LV myocardium has previously been characterized,^5^ the sexual dimorphism of ischemic heart disease, and a paucity of sex-specific investigation to date,^4^ suggests sex-specific transcriptional responses might explain these mechanisms. In this investigation, we identified sex-specific changes in gene expression in the human LV myocardium post-ischemia. Our whole-genome RNA-seq analysis identified sex differences in the transcriptional response to ischemia and functional analysis of these genes revealed enrichment of several biological processes. An eQTL analysis identified sex differences in the genetic regulation of gene expression post-ischemia, and cell-type deconvolution analysis showed sex-bias in proportion of specific cell types.

### Relative up-regulation in females

*FAM5C* is expressed in coronary artery smooth muscle and endothelium where over-expression increases monocyte adhesion.^31^ It has been previously demonstrated that elevated inflammatory biomarkers are better predictors of outcomes in ischemic heart disease in women than are traditional biomarkers,^31^ suggesting that women experience a larger cardiac inflammatory response than do men following myocardial ischemia. The greater up-regulation of *FAM5C* in women in response to ischemia in our study indicates that women may experience increased monocyte adhesion, and therefore a larger inflammatory infiltrate, in response to ischemia than do men. FAM5C may therefore serve as a novel target to modulate cardiac inflammation following ischemia.

The superfamily of phospholipase A_2_ enzymes is composed of 5 subgroups, several of which are biomarkers for ischemic events.^32^ It has recently been implicated in endocytosis^33^ and in the biosynthesis of a class of biological molecules called *N*-acyl phosphatidylethanolamines (NAPEs).^34^ cPLA_2_E has been observed to share an endosomal binding site with PLD2.^33^ Phospholipase D2 localizes to the sarcolemmal membrane and regulates endocytosis of angiotensin-II type I receptor, which is injurious to the myocardium after ischemia/reperfusion injury.^35, 36^ In our study, *PLD2* was significantly up-regulated in both sexes, but to a lesser degree in women. Changes in *PLA2G4E* expression, and its relationship to *PLD2*, may therefore modulate cardiac angiotensin action. Additionally, *PLA2G4E* transfected cells accumulated downstream NAPE metabolites, which may come from a specific source of NAPEs^34^ and would reveal an injurious modulation of a specific, potentially harmful reservoir of biological molecules in only females during myocardial ischemia.

*CYP1A1*, which was down-regulated in males and up-regulated in females in response to ischemia, metabolizes estrogens in the myocardium to form a 2-hydroxy metabolite (2H),^37, 38^ which can then be metabolized to a 2-methoxy product (2ME).^39^ 2ME has been demonstrated to be anti-angiogenic and augment apoptosis in endothelial cells,^40^ and, in the myocardium, both metabolites increase apoptosis in ischemia-reperfusion injury.^41, 42^ Furthermore, *ESR1*, which encodes estrogen receptor alpha (ERα), was expressed similarly between the sexes at baseline, but showed a differential response to ischemia with up-regulation in males and down-regulation in females in response to ischemia. ERα has been shown to protect against ischemic injury.^25^ Estrogen is an established cardioprotective agent in pre-menopausal females, and hormone replacement therapy (HRT) has been investigated in the reduction of cardiovascular events. However, HRT has been found to increase cardiovascular disease in postmenopausal females within the first several years of treatment,^44, 45^ though no patients in this study were undergoing HRT. Our results suggest that the changes seen in post-menopausal females, expression of genes that increase estrogen metabolism and reduce estrogen binding, may be injurious in ischemic heart disease, and/or that the increased action of estrogen in males following ischemia may be a protective mechanism not shared by post-menopausal females.

### Relative down-regulation in females

D3, down-regulated to a significantly greater degree in females, is one of three iodothyronine deiodinases, and is responsible for deactivating thyroid hormone.^46^ Myocardial *DIO3* induction has been observed in infarcted rat models.^47^ While in both sexes there were changes in iodothyronine deiodinase 2 and 3 gene expression, thyroid hormone levels were not measured. Despite a cardioprotective indication for thyroid hormone post ischemia,^48^ it may not be metabolized in the hearts of patients undergoing cardiac surgery.^49^ Epigenetic down-regulation of *DIO3*, which has been observed in atherosclerotic arteries,^50^ may therefore be responsible for the detected change in females. As atherosclerosis is an inflammatory disease,^51^ down-regulation of *DIO3* may reflect a sex-specific injurious inflammatory response.

Chymase is the main angiotensin II-forming enzyme in the human heart,^56^ and its inhibition has been shown to be cardioprotective following ischemia/reperfusion injury and myocardial infarction.^57^ However, chymase activity from cardiac mast cells has also been implicated in angiogenesis in the myocardial microvasculature.^58^ The down-regulation of *CMA1* in females and no change in males due to ischemia might elucidate why the cardiac microvasculature has been shown to contribute to the progression of ischemic heart disease to a greater degree in females.^59^

### cis-eQTL

The effect on gene expression of the SNP-by-sex interaction response QTLs identified in this investigation have not been previously described. We demonstrated, that not only could *cis*-eQTLs have sex-dependent effects on gene expression at baseline, but also in response to ischemia. One such gene to demonstrate such an interaction is VGLL4. VGLL4 binds to TEAD4 and forms a critical transcription factor complex that regulate vascular endothelial growth factor A (VEGFA), a well-known antigenic cytokine which gets activated in response to ischemia to restore blood flow.^60^ rs11707097-AA genotype is associated with decreased VGLL4 expression in post-ischemia in females only, suggesting that the genotype determines its sex-specific down-regulation due to ischemia. While the eQTLs identified here may indicate that women with certain genotypes are at an increased risk of ischemic injury, some genes had few homozygous genotypes. Nevertheless, SNP-by-sex interaction response QTLs clearly warrant further investigation.

### Limitations

This investigation is unique in several ways. Sample collection was performed at the same physical location on the LV apex by the same surgeon at the same time in the operation, thereby reducing sample variability. The time from sample collection to RNA preservative was about 1 minute, resulting in extremely high-quality RNA, and, whereas similar studies often obtain tissue post-mortem or from cardiac transplantation, tissue was collected from live, non-failing hearts. While the sampled hearts had concentric hypertrophy from long-standing aortic stenosis, all samples were collected during non-urgent, elective procedures, and risk factors were optimally medically managed beforehand.

Despite the strengths of this investigation, there remain limitations. Ischemia induced by cold cardioplegia is not equivalent to *in vivo* ischemia, akin to coronary artery ligation in rodents. The results of this experiment may therefore underestimate the extent of ischemic injury.

Additionally, down-regulation of gene expression may be due to cold temperature. However, the release of cardiac-specific biomarkers to levels that would qualify as a myocardial infarction in the non-surgical population suggest ischemic changes in response to aortic cross-clamping and cardiopulmonary bypass. Lastly, while we were not able to distinguish individual cell-types using single-cell sequencing, and the small amount of tissue collected prohibited protein-level validation, our cell-type enrichment analysis demonstrates that intra-individual samples exhibit similar cell type composition profiles that are unaffected by ischemia.

### Conclusion

We have demonstrated that the human LV demonstrates significant sex-specific changes in gene expression as a result of cold cardioplegia-induced ischemia during CPB. The results of this investigation provide a valuable foundation for examining the sexual dimorphism of surgical outcomes in CPB, and may, by extension, help elucidate the mechanisms underlying sex differences in ischemic heart disease. The genes, pathways, and variants revealed by this experiment should serve as targets for further exploration, and, more generally, support the necessity of sex-specific experimentation in the advancement of human health and precision medicine for all.

## Acknowledgements

We acknowledge the contribution to sample collection by the Perioperative Genomics Center and its staff: James Gosnell, RN; Kujtim Bodinaku, MD; Svetlana Gorbatov, MPH. We also thank James Bell for graphical assistance.

## Online Data Supplements

### Figures

Supplemental Figure 1: Effect of filtering on stabilizing mean-variance trend

Supplemental Figure 2: Principal component clustering of gene expression profiles of 220 heart samples shows clustering by A) individual and by B) Pre- and Post-ischemia.

Supplemental Figure 3: Cell-type enrichment analysis of TRANSCRIBE left ventricular samples and GTEx samples

Supplemental Figure 4: Stratification of enrichment score by sex and baseline and ischemia conditions

### Tables

Supplemental Table 1: Complete lists of significantly differentially expressed genes in each sex (BH adjusted p-value < 0.05)

Supplemental Table 2: Complete list of genes with sex-specific differential expression due to ischemia (BH adjusted p-value < 0.05)

Supplemental Table 3: Complete functional analysis results

Supplemental Table 4: Complete *cis*-eQTL results

